# The Regulation of Nucleic Acid Vaccine Responses by the Microbiome

**DOI:** 10.1101/2023.02.18.529093

**Authors:** Andrew M. F. Johnson, Kevin Hager, Mohamad-Gabriel Alameh, Phuong Van, Nicole Potchen, Koshlan Mayer-Blackwell, Andrew Fiore-Gartland, Samuel Minot, Paulo J. C. Lin, Ying K. Tam, Drew Weissman, James G. Kublin

## Abstract

**Summary:** Nucleic acid vaccines, including both RNA and DNA platforms, are key technologies that have considerable promise in combating both infectious disease and cancer. However, little is known about the extrinsic factors that regulate nucleic acid vaccine responses and which may determine their effectiveness. The microbiome is recognized as a significant regulator of immune development and response, whose role in regulating some traditional vaccine platforms has recently been discovered. Using germ-free and specific-pathogen-free mouse models in combination with different protein, DNA, and mRNA vaccine regimens, we demonstrate that the microbiome is a significant regulator of nucleic acid vaccine immunogenicity. While the presence of the microbiome enhances CD8+ T cell responses to mRNA lipid nanoparticle (LNP) immunization, the microbiome suppresses immunoglobulin and CD4+ T cell responses to DNA-prime, DNA-protein-boost immunization, indicating contrasting roles for the microbiome in the regulation of these different nucleic acid vaccine platforms. In the case of mRNA-LNP vaccination, germ-free mice display reduced dendritic cell/macrophage activation that may underlie the deficient vaccine response. Our study identifies the microbiome as a relevant determinant of nucleic acid vaccine response with implications for their continued therapeutic development and deployment.

## Introduction

Vaccines are among the most effective tools for preventing human morbidity and mortality from infectious disease. Recently, nucleic acid vaccines encompassing both RNA and DNA platforms have emerged as a key technology offering potential advances over traditional protein-adjuvant approaches.^1–3^ In particular, nucleic acid vaccines promote enhanced cellular as well as humoral immunity, which may be particularly effective in combating intracellular pathogens and cancers.^4,5^ Furthermore, advantages in the speed and flexibility of design and production make nucleic acid vaccines particularly valuable in combating emerging pathogens.^1,6^ RNA-based COVID-19 vaccines combining modified mRNA molecules with lipid nanoparticle (LNP) technology were brought to the clinic within a year of SARS-CoV-2 emerging in humans and their remarkable clinical efficacy has played a vital role in controlling the COVID-19 pandemic.^7,8^ DNA vaccines demonstrated impressive preclinical results, however this has so far not translated easily to the human setting where DNA vaccine immunogenicity has been lower than was anticipated.^9^ Modified DNA vaccine delivery regimens have been developed to overcome low immunogenicity, including combining DNA and protein vaccines in prime-boost regimens that enhance the quality and quantity of the elicited responses relative to either modality alone.^10–14^

Despite considerable past success, vaccine effectiveness remains hard to predict and considerable inter-individual heterogeneity in the immune response to vaccines can limit their clinical effectiveness.^15,16^ The determinants of vaccine response across individuals remain poorly understood, especially for nucleic acid vaccines, and likely involve a variety of immune-modulatory factors, such as age, genetics, nutrition, past microbial infections, and the microbiome.^16–18^ The microbiome is an important regulator of immune development and response.^19,20^ Data from preclinical mouse models have demonstrated that components of the microbiome can influence the immunogenicity of several vaccine platforms,^21^ including trivalent inactivated influenza vaccine,^22^ canonical infant vaccines,^23^ adenoviral *Mycobacterium tuberculosis* vaccine,^24^ live-attenuated rabies vaccine,^25^ and proteins adjuvanted with complete Freud’s adjuvant,^26^ Alum,^27^ CpG,^28^ or cholera toxin.^29^ Associations between vaccine response and microbiota composition have also been observed in human cohorts,^30–35^ and microbiota-disrupting antibiotic treatment of human subjects has been associated with altered influenza and rotavirus vaccine immunogenicity.^33,36^ However, whether the microbiome regulates RNA or DNA vaccine immunogenicity remains undescribed. Furthermore, the precise nature of microbiome control over existing vaccines remains to be resolved with contrasting effects reported depending on factors such as age of mice or route of administration,^23,29^ and for similar experiments reproduced across institutions.^22,23^

We explored the contribution of the microbiome to different vaccine technologies by comparing the immune responses of germ-free (GF) and specific-pathogen-free (SPF) mice immunized with different protein-adjuvant vaccine regimens, a DNA-prime DNA-protein-boost vaccine regimen,^10,37^ and a single dose nucleoside-modified mRNA-lipid nanoparticle (LNP) vaccine.^38–40^ Our data indicate contrasting roles for the microbiome in regulating protein, DNA-prime DNA-protein-boost, and mRNA-LNP vaccine responses and provide a basis for further interrogation of the microbiome-dependent mechanisms that regulate nucleic acid vaccine immunogenicity.

## Results

### Humoral and cellular responses to protein-adjuvant vaccines in GF and SPF mice

GF and SPF C57Bl/6J mice were immunized subcutaneously (s.c.) on day 0 and day 21 with Ovalbumin (OVA) protein formulated with either Alhydrogel® (OVA/Alum), Addavax (OVA/Addavax), or AS01 (OVA/AS01) adjuvants. GF and SPF mice injected with OVA in phosphate-buffered saline (OVA/PBS), or PBS alone (Naïve) were employed as negative controls. Three weeks after the final immunization anti-OVA IgG1, IgG2c, and IgG2b were quantified in serum by ELISA (**Figure 1A**). Anti-OVA-IgG1, -IgG2c, and -IgG2b titers were all highest in OVA/AS01-immunized mice, followed by OVA/Alum- and OVA/Addavax-immunized mice. This effect was strongest for anti-OVA IgG2c consistent with enhanced type-1 polarization by AS01 in mice compared to Alum or Addavax. There were no significant differences in anti-OVA IgG1, -IgG2c or -IgG2b between vaccine-matched GF and SPF mice indicating a minimal contribution of the SPF microbiome to the humoral responses of these protein-adjuvant vaccines. Mice receiving OVA/PBS displayed anti-OVA-IgG1, -IgG2c and -IgG2b titers 4-5 log units lower than adjuvanted mice as expected. Anti-OVA-IgG2b titers were slightly, but significantly higher in OVA/PBS-injected SPF mice than OVA/PBS-injected GF mice consistent with a role of the SPF microbiome in supporting T-independent humoral responses to unadjuvanted antigen.

**Figure 1:**
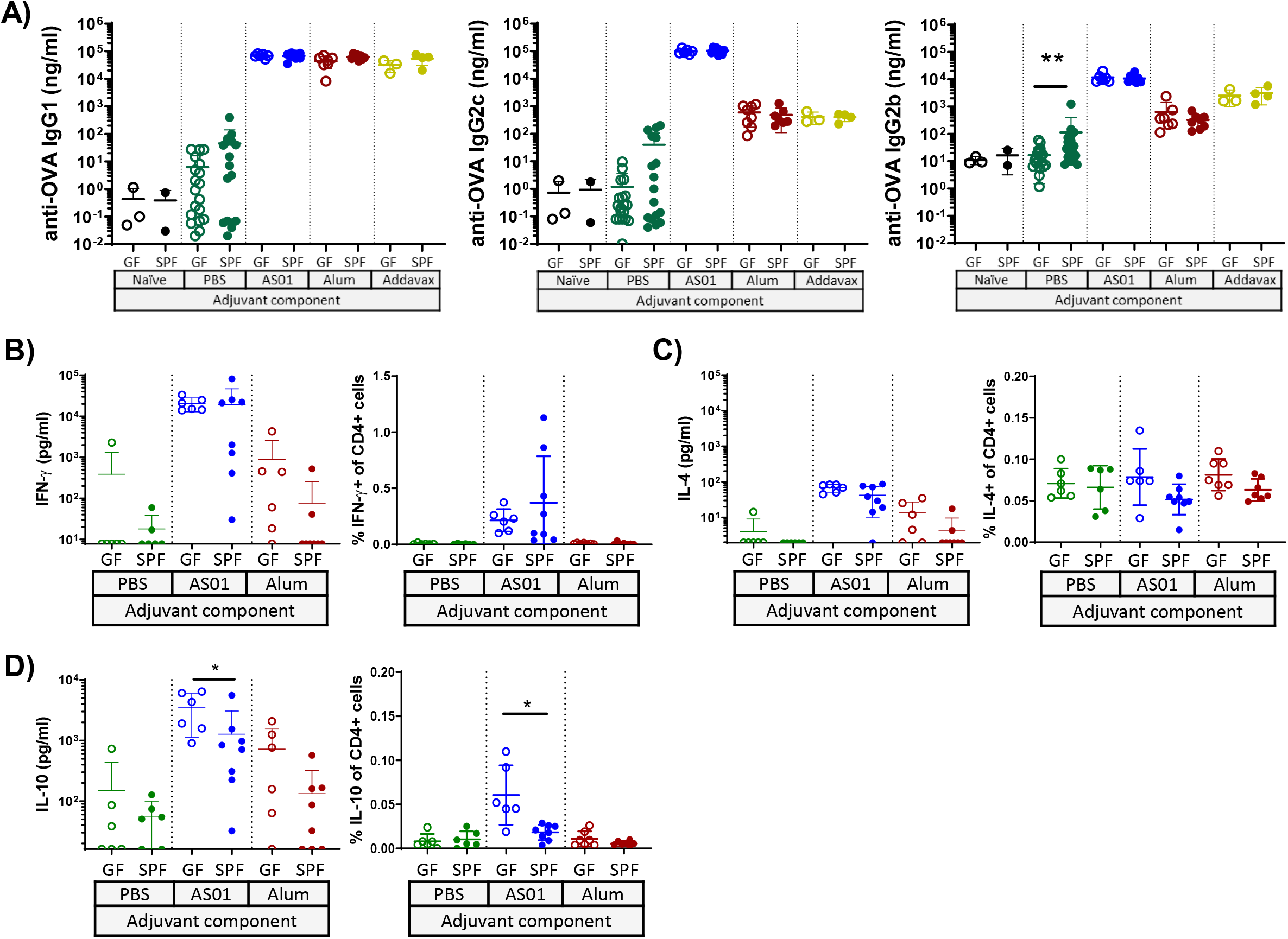
Humoral and cellular responses to OVA immunization In GF and SPF mice. **(A)** OVA-specific IgG1, IgG2c, IgG2b titers in serum and OVA-stimulated splenocyte **(B)** IFN-γ, **(C)** IL-4 and **(D)** IL-10 responses from GF or SPF mice either unimmunized (naïve) (n = 3), or immunized with with OVA in PBS (PBS) (n = 17-19), OVA with AS01 (AS01) (n = 6-8), OVA with Alum (Alum) (n = 7-8), or OVA with Addavax (Addavax) (n = 3-4). Samples falling below level of cytokine detection (LOD) by ELISA were set at half the LOD, which is the beginning of y-axis, for visualization. Each data-point represents an individual mouse from one experiment. Vaccine-matched GF and SPF groups were compared for statistical significance by Mann Whitney test, ** p < 0.01.

In order to assess vaccine-induced T cell responses we re-stimulated splenocytes from OVA/AS01-, OVA/Alum-, and OVA/PBS-immunized mice with OVA protein *ex vivo.* T cell responses were quantified both by intracellular cytokine staining (ICS) flow cytometry and by the concentration of cytokine released into the culture supernatant relative to unstimulated controls **(Figure 1B-D)**. OVA-stimulated IFN-γ+ CD4+ T cell proportions and IFN-γ concentration in the culture supernatant were markedly increased in OVA-stimulated splenocyte cultures from OVA/AS01-immunized mice compared to OVA/Alum- or OVA/PBS-immunized mice consistent with the enhanced Type-1 polarization activity of AS01 **(Figure 1B)**. OVA-stimulated IL-4 concentration was also increased in splenocyte cultures from OVA/AS01-, but not OVA/Alum-immunized mice, compared to OVA/PBS controls, although these IL-4 responses were low (less than 100 pg/ml) and were not detectable by ICS. IFN-γ and IL-4 concentrations were not significantly different in OVA-stimulated splenocyte cultures from vaccine-matched GF and SPF mice **(Figure 1B-C)**. Finally, both IL-10 cytokine concentration and the proportion of IL-10+ CD4+ T cells were increased in OVA-stimulated splenocyte cultures from GF OVA/AS01-immunized mice and these IL-10 responses were significantly higher than SPF OVA/AS01-immunized mice (**Figure 1D)**, suggesting that the microbiome may play a role in suppressing IL-10 responses in OVA/AS01-immunized mice.

### Humoral responses to protein-adjuvant vaccines in selectively colonized mice

In order to extend our assessment of microbiome influence on protein-adjuvant immunization beyond the conventional SPF microbiome of C57Bl6/J mice, we employed two distinct colonization approaches. Firstly, we infected GF mice with a five-member microbial consortia (PRO-consortia) consisting of *Akkermansia mucinophilia, Prevotella copri, Fusobactrium varium, Colinsella aerofaciens,* and Segmented Filamentous Bacteria (SFB) selected for their reported immune-stimulatory activities *in vivo*.^41–43^ GF mice were infected by oral gavage beginning 7 days prior to immunization with OVA/Addavax in a prime-boost regimen. Quantification of bacterial 16s rDNA copy number in the stool of colonized mice with species specific primers indicated that *Prevotella copri* failed to colonize, while the remaining four-members of the consortia displayed stable colonization levels over the course of the experiment (**Supplementary Figure 1A**). There was no difference in OVA-specific IgG titers between PRO-consortia-colonized mice and uncolonized GF controls (**Supplementary Figure 1B**). Secondly, we infected GF mice with cryopreserved cecal material obtained from WILD-re-derived (WILD-R) mice. GF mice were orally gavaged on three consecutive days with either WILD-R or SPF material or remained uncolonized (GF), while standard SPF-raised mice (SPF-R) were employed as controls. Two weeks after colonization blood samples were collected from GF, WILD-R-colonized, SPF-colonized (SPF-C), and SPF-R mice and the proportions of circulating immune cells measured by flow cytometry with no significant differences detected between groups (**Supplementary Figure 2A**) suggesting that no sustained inflammatory reaction was taking place as a result of exposure to WILD-R or SPF material. Mice were immunized four weeks post-colonization with OVA/Alum in a prime-boost regimen. No differences between OVA-specific IgG1, IgG2b or IgG2c were detected between GF, SPF-colonized or WILD-R-colonized mice (**Supplementary Figure 2B**). 16S rDNA sequencing of cecal contents of WILD-R and SPF-C mice indicated substantial differences in the microbiome communities between WILD-R and SPF-C mice as shown by principal coordinate analysis using Bray-Curtis distance metric at the species level (**Supplementary Figure 2C**). Several families of bacteria were differentially abundant between WILD-R-colonized and SPF-C mice with *Bifidobacteriaceae, Desulfovibrionaceae, Eggerthellaceae, Marinilabiliaceae, Ordoribacteraceae, Sphingobacteriaceae, Streptococcaceae, Tannerllaceae* and the candidate phylum *Candidatus saccharibacteria* significantly increased in WILD-R-colonized mice, and *Bacteroidaceae, Clostridiaceae, Eubacteriaceae and Valitaleaceae* were significantly increased in SPF-C mice (**Supplementary Figure 2D**). The alpha diversity, as measured by Simpson or Shannon indices, of WILD-R-colonized mice trended towards being increased compared to SPF-C mice (**Supplementary Figure 2E**). However, the alpha diversities of WILD-R-colonized mice were notably lower than the donor WILD-R material, indicating incomplete transplant of WILD-R microbiome diversity by this procedure. Indeed, the phyla most differentially abundant between donor WILD-R and SPF material, Proteobacteria consisting mostly of *Helicobacter* species, failed to colonize GF mice indicating the challenges using this transplant approach to fully recapitulate the WILD-R microbiome community (**Supplementary Figure 2F**). Collectively, these data show that addition of select immunostimulatory species or reconstitution with a WILD-R microbiome community were insufficient to modulate OVA/Addavax or OVA/Alum vaccine immunogenicity in mice.

### Humoral and cellular responses to DNA-prime, DNA-protein-boost immunization in GF and SPF mice

The vaccine DNA-HIV-PT123 consists of three plasmids encoding HIV gp140 env, gag, and pol-nef fusion proteins and has been employed in several clinical studies.^44,45^ We employed DNA-HIV-PT123 in GF and SPF mice using a DNA-HIV-PT123-prime, DNA-HIV-PT123 + MF59-adjuvanted gp120 protein-boost regimen that mirrors the regimen employed in HIV Vaccine Trial Network (HVTN) 108 and 111 clinical trials (**Figure 2A**). Mice were immunized subcutaneously by needle-syringe on day 0 with DNA-HIV-PT123 alone, and on day 14 and day 35 with both DNA-HIV-PT123 and gp120 protein adjuvanted with MF59 (**Figure 2A**). Three weeks after the final immunization anti-gp120 IgG was quantified in serum by ELISA, and splenocytes were re-stimulated with gp120 protein to measure T cell responses by ICS flow cytometry and by the concentration of cytokine released into the supernatant relative to unstimulated controls. GF mice receiving this immunization regimen displayed significantly higher mean anti-gp120 IgG antibody titers than SPF mice (**Figure 2B**). A post-hoc Levene’s test also indicated that anti-gp120 IgG titers of GF mice were significantly less variable than those of SPF mice (p = 0.0102). GF mice displayed significantly higher IFN-γ+ and TNF-α+ CD4+ T cell responses than SPF mice measured by ICS (**Figure 2C**) following gp120 splenocyte re-stimulation. Consistent with this, IFN-γ concentrations were also increased in the supernatant of gp120-restimulated splenocyte cultures from GF compared to SPF mice (**Figure 2D**). These data indicate that the microbiome suppresses both humoral and cellular immunogenicity to this DNA-HIV-PT123-prime, DNA-HIV-PT123 + gp120/MF59-boost regimen.

**Figure 2:**
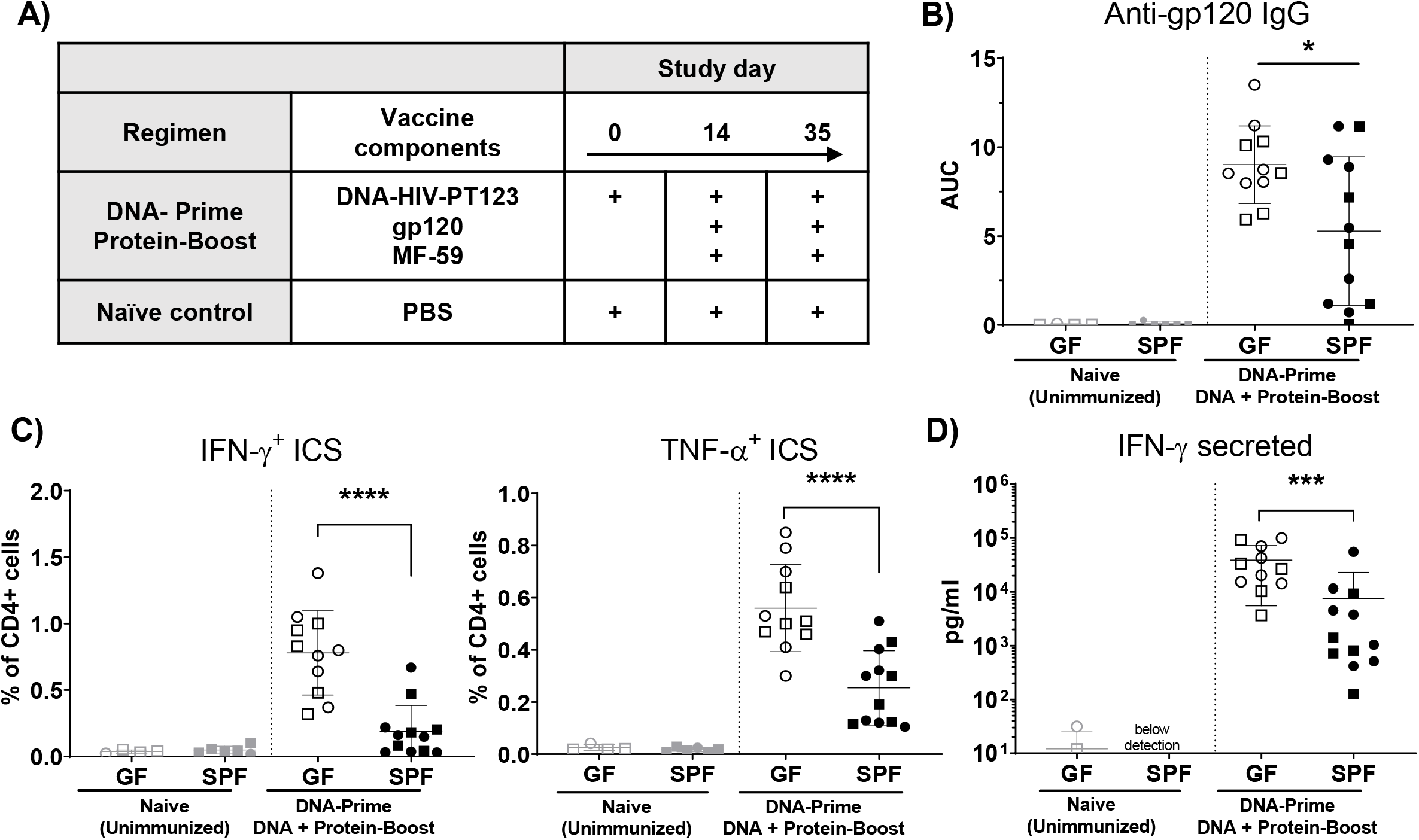
Humoral and cellular responses to DNA-HIV-PT123-prime, DNA-HIV-PT123 + gp120 protein-boost vaccination in GF and SPF mice. **(A)** Summary of DNA-HIV-PT123 and gp120/MF59 vaccine regimen. **(B)** Serum gp120-specific IgG titers and **(C)** proportions of IFN-γ+ and TNF-α+ CD4+ T cells and **(D)** IFN-γ concentration in supernatant of gp120-stimulated splenocyte cultures from immunized (DNA+gp120+MF59) (n = 11-12) and unimmunized (naïve) (n = 4-5) GF and SPF mice. Samples falling below level of IFN-γ detection (15.6 pg/ml) were set at 7.81 pg/ml for visualization. Each data-point represents an individual mouse from two experiments that gave very similar results; cohort 1 (circles), cohort 2 (squares). * p < 0.05 (Welch’s T test), **** p < 0.0001 and *** p < 0.001 (Mann-Whitney test).

In order to define more precisely which components of this DNA-HIV-PT123-prime, DNA-HIV-PT123 + gp120/MF59-boost regimen the microbiota regulates, we compared the humoral and cellular responses of GF and SPF mice immunized with either (1) DNA-HIV-PT123 alone, (2) DNA-HIV-PT123 with MF59, (3) DNA-HIV-PT123 with unadjuvanted gp120, (4) gp120 with MF59, or (5) gp120 alone. No significant differences were observed in serum anti-gp120 IgG titers, gp120-responsive IFN-γ+ and TNF-α+ splenic CD4+ T cells, or gp120-induced splenocyte IFN-γ secretion between vaccine-matched GF and SPF mice for any of these partial regimens (**Figure 3**), indicating that the full DNA-HIV-PT123 prime, DNA-HIV-PT123 + gp120/MF59 boost regimen is necessary to reveal microbiome-dependent regulation in this model. Notably, anti-gp120 IgG and CD4+ T cell ICS responses were generally low in mice receiving DNA-HIV-PT123 alone, DNA-HIV-PT123 with MF59, or DNA-HIV-PT123 with unadjuvanted gp120 protein with some of the mice undergoing these regimens failing to mount a response above background even after three 100 μg doses of DNA-HIV-PT123 vaccine (**Figure 3**). Serum anti-gag IgG titers were also not detectable in nearly all DNA-HIV-PT123-immunized mice despite plasmids encoding the Gag immunogen being present (**Supplementary Figure 3**). This indicates that, in the absence of subsequent protein-adjuvant boost, s.c. needle-syringe delivery of DNA-HIV-PT123 is poorly immunogenic in mice regardless of gnotobiotic condition, thus limiting our ability to assess the contribution of the microbiome to regulation of DNA-HIV-PT123 immunization alone in this model.

**Figure 3:**
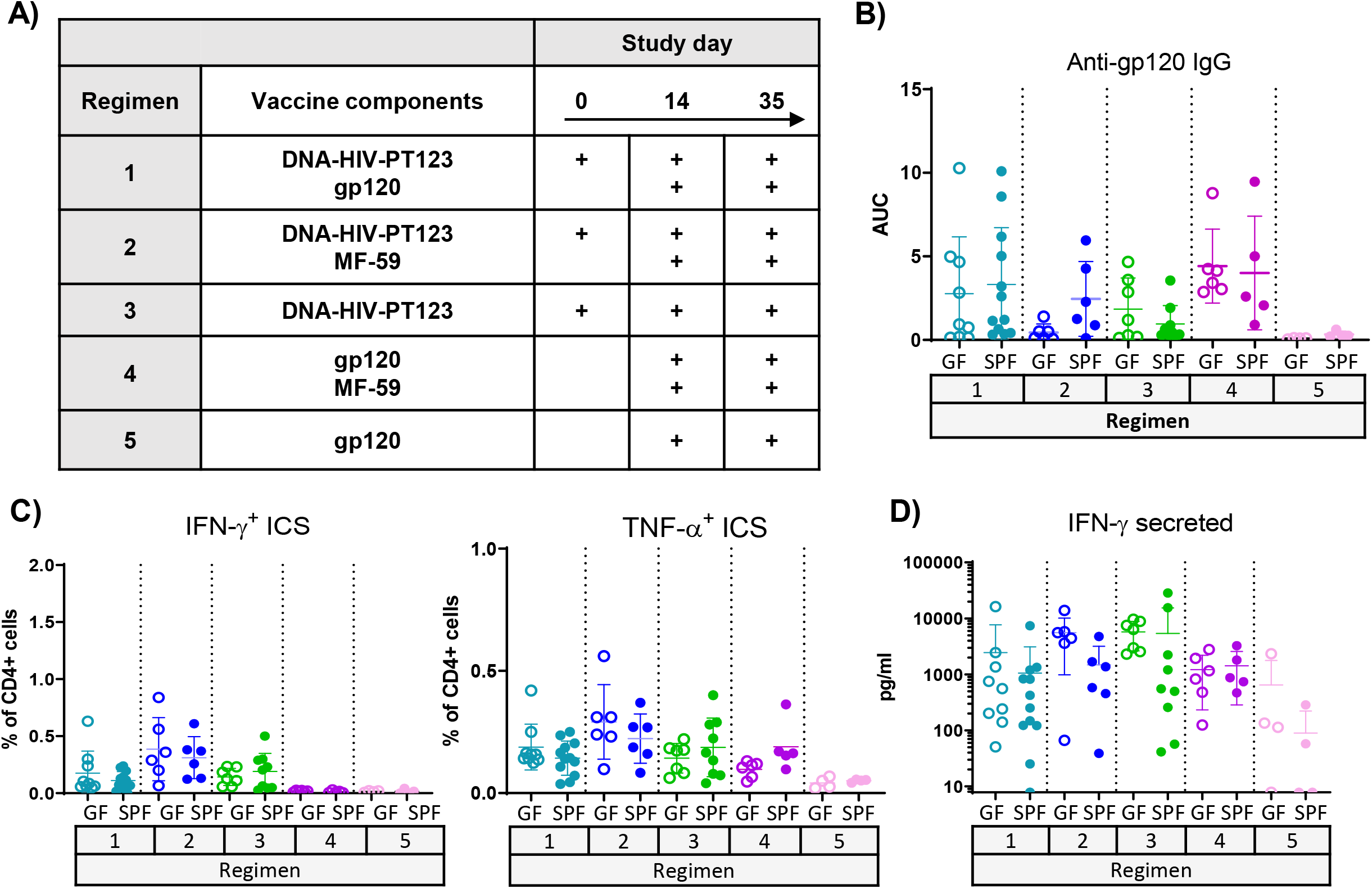
Humoral and cellular responses to partial DNA-HIV-PT123 and gp120 vaccine regimens in GF and SPF mice. **(A)** Summary of DNA-HIV-PT123, gp120 MF59 vaccine regimens employed. **(B)** Serum gp120-specific IgG titers and **(C)** proportions of IFN-γ+ and TNF-α+ CD4+ T cells and **(D)** IFN-γ concentration in supernatant of gp120-stimulated splenocyte cultures from immunized (DNA+gp120+MF59) (n = 11-12) and unimmunized (naïve) (n = 4-5) GF and SPF mice. Samples falling below level of IFN-γ detection (15.6 pg/ml) were set at 7.81 pg/ml for visualization. Each data-point represents an individual mouse from one or two experiments depending on group. There were no significant differences between vaccine-matched GF and SPF groups (Mann-Whitney test).

### Humoral and cellular responses to mRNA-LNP immunization in GF and SPF mice

To investigate whether the microbiome contributes to the immunogenicity of a nucleoside-modified mRNA-LNP vaccine, as used in Moderna and Pfizer/BioNTech COVID-19 vaccines, we immunized GF and SPF mice subcutaneously with a single immunization of mRNA-LNP vaccine encoding SARS-CoV-2 Spike protein; mice were euthanized 14 days later. Anti-Spike IgG titers were determined in serum and CD4+ and CD8+ T cell responses were quantified by ICS flow cytometry following splenocyte re-stimulation with a spike peptide pool. There were no significant differences in anti-Spike IgG titers or in the proportions of CD4+ T cells producing IFN-γ, IL-2, IL-4, IL-5, TNF-α, or IL-17 between GF or SPF mRNA-LNP-immunized mice (**Figure 4A and 4B**). However, there were significant reductions in the proportions of CD8+ T cells producing IFN-γ, IL-2 or TNF-α, and co-expressing IFN-γ+ and CD107α+ in GF compared to SPF mice following mRNA-LNP immunization (**Figure 4C**). These data indicate that CD8+ T cell responses to the mRNA-LNP immunization utilized in this study are enhanced by the presence of the microbiome.

**Figure 4:**
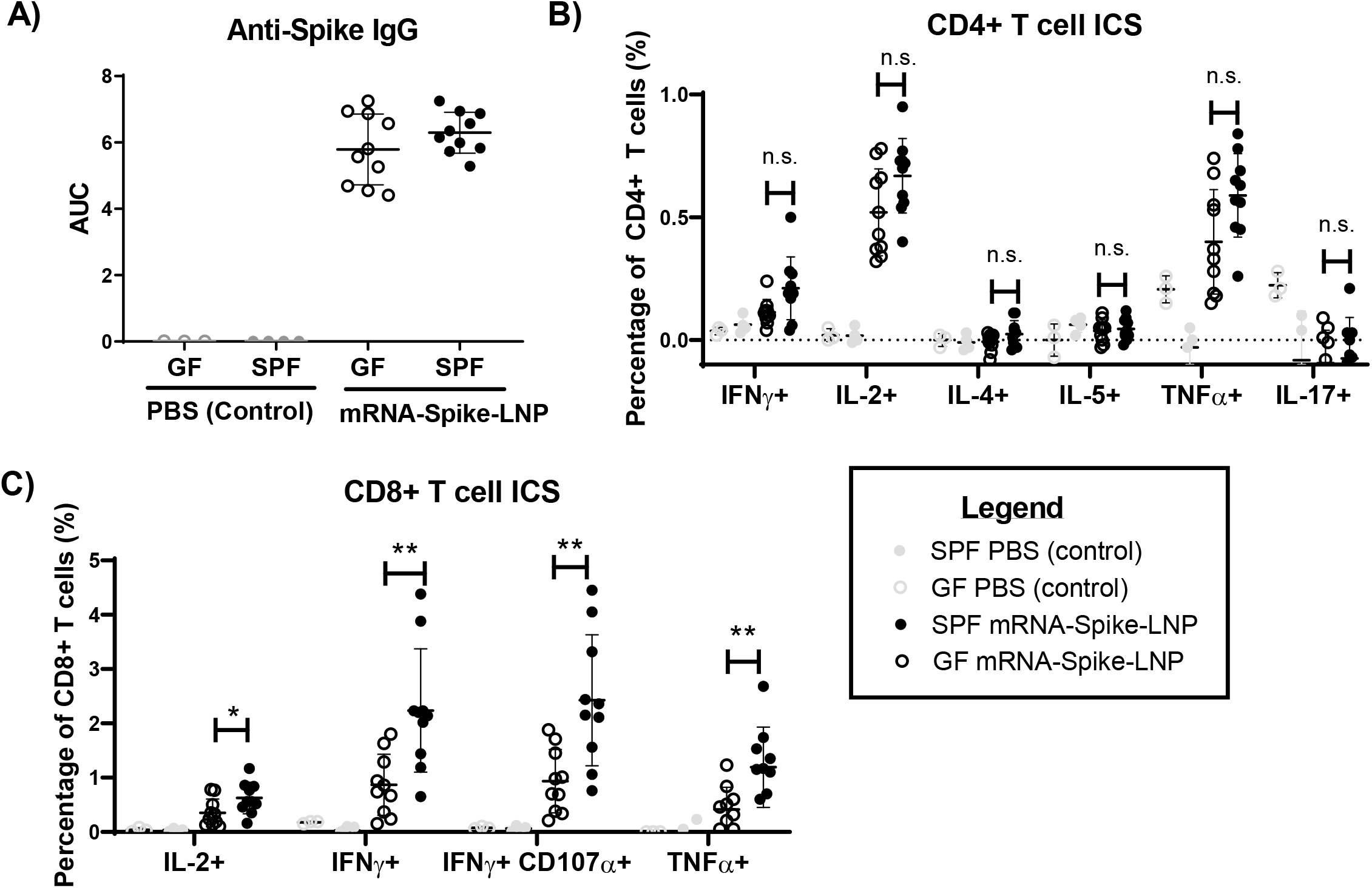
Humoral and cellular Responses to mRNA-LNP immunization in GF and SPF mice. **(A)** Serum Spike-specific IgG **(B)** Spike peptide pool-stimulated CD4+ T cell cytokine responses quantified by ICS, and **(C)** Spike peptide pool-stimulated CD8+ T cell cytokine responses quantified by ICS in unimmunized (PBS Control) and mRNA-Spike-LNP-immunized GF (n = 10) and SPF (n = 12) mice from one experiment. * *p* < 0.05 and ** *p* < 0,01 (Mann-Whitney test).

To provide insight into the potential mechanisms by which the microbiome contributes to mRNA-LNP immunization, we completed cellular immune profiling of the early innate response to mRNA-LNP 24 hours post-immunization in spleen tissue of GF and SPF mice. There was a dramatic activation of CD11b+ Ly6Chi monocytes following mRNA-LNP-immunization, with nearly all Ly6Chi monocytes in the spleens of immunized SPF mice displaying upregulation of the dendritic cell (DC)/macrophage marker, CD11c (**Figure 5A**). Interestingly the total proportion of Ly6Chi monocytes, as well as the proportion of Ly6Chi monocytes expressing CD11c was reduced in GF compared to SPF mRNA-LNP-immunized mice (**Figure 5A**) indicative of reduced Ly6Chi monocyte activation in GF animals. We employed a gating strategy to assess neutrophil and DC/macrophage populations based on expression of CD11b, Ly6G, CD64, Ly6C, CD11c and MHC-II (**Figure 5B**). mRNA-LNP immunization resulted in an accumulation of neutrophils (CD11b+Ly6Ghi), which was reduced in GF mice (**Figure 5C**). We focused our initial analysis on distinct populations of Ly6C+ and CD64+ DC/macrophage cells, which have previously been associated with inflammation and the response to vaccination,^46,47^ and Ly6C-CD64-conventional DCs defined as follows: Ly6C+CD11c+MHC-II+ cells (Ly6C+ DCs), CD64+ Ly6C-/lo CD11c+ MHC-II+ cells (CD64+Ly6C-/lo DC/MP), and Ly6C-CD64-CD11c+MHC-II+ cells (CD64-Ly6C-cDCs) (**Figure 5B)**. The expression of other DC/macrophage-associated cell surface proteins, such as CD11b, CD80, and CD86, on these sub-populations is shown in **Supplementary Figure 4**. The proportion of Ly6C+ DCs in the spleen was increased by mRNA-LNP immunization in both SPF and GF animals (**Figure 5C**). Interestingly, there was also an increase in the proportion of CD64+Ly6C-/lo DC/MPs following mRNA-LNP immunization in SPF mice that was not evident in GF mice (**Figure 5C**). Meanwhile, CD64-Ly6C-cDCs, including XCR1+, CD8a+, and CD11b+ DCs were unchanged by mRNA-LNP immunization in either SPF or GF settings (**Figure 5C** and **Supplementary Figure 5**). Collectively, these data demonstrate the significant innate immune activation which occurs following mRNA-LNP immunization and highlight differences in both monocyte activation and CD64+Ly6C-/lo DC/macrophage accumulation between SPF and GF mice.

**Figure 5:**
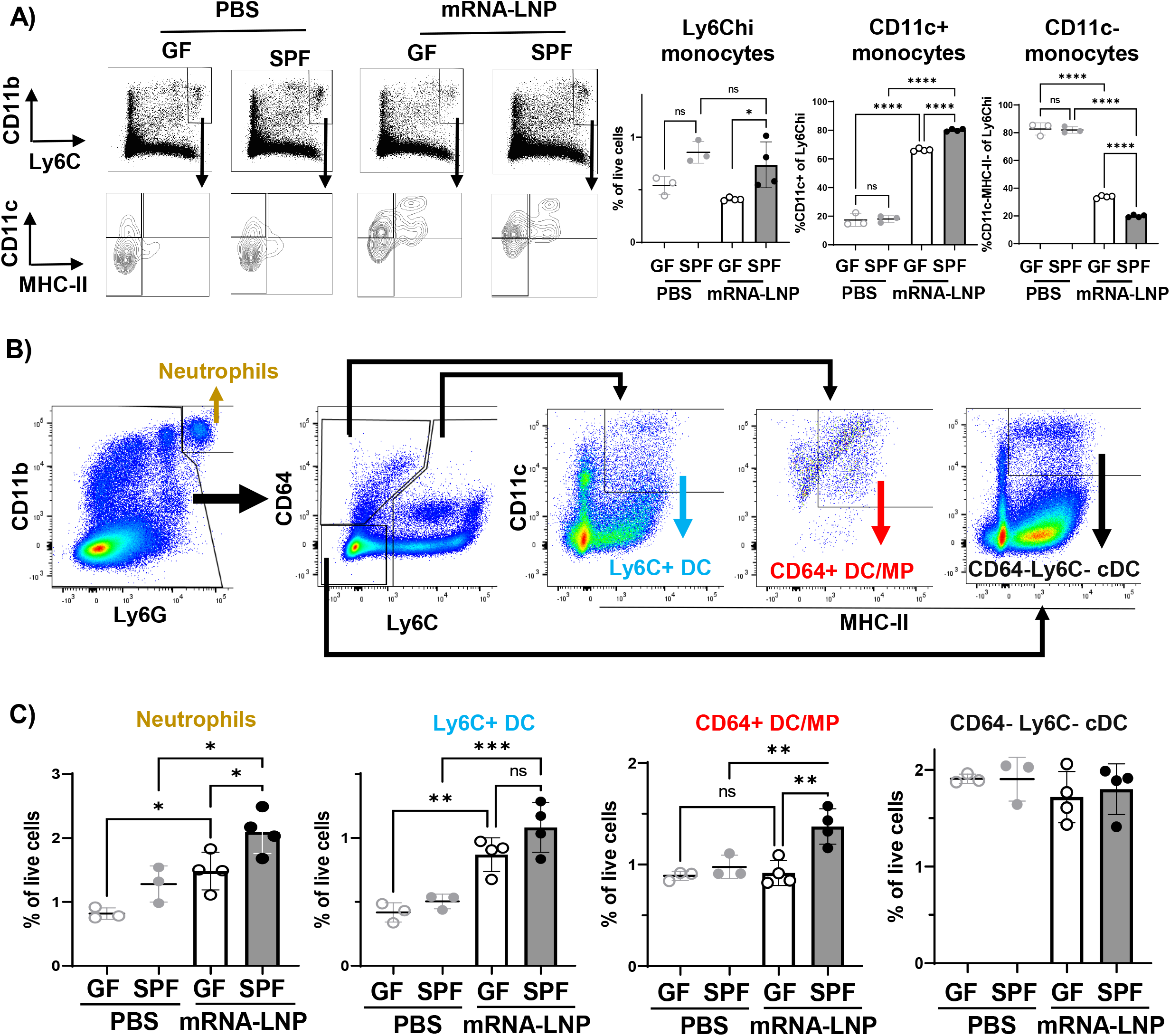
Flow cytometry profiling of innate immune response to mRNA-LNP in GF and SPF mice. **(A)** Representative flow cytometry and graphical summaries of Ly6Chi monocyte gating and monocyte CD11c vs MHC-II expression in PBS-treated or mRNA-LNP-immunized GF and SPF mice. **(B)** Gating strategy to identify Neutrophils and Ly6C+, CD64+ and CD64-Ly6C-DC/macrophage populations amongst splenocytes from PBS-treated or mRNA-LNP-immunized GF and SPF mice. **(C)** Graphical summaries of Neutrophil and Ly6C+, CD64+ and CD64-Ly6C-DC/macrophage proportions in PBS-treated or mRNA-LNP-immunized mice. Each data-point on graphs represents a single mouse (n = 3 per PBS group, n = 4 per mRNA-LNP group from one experiment). * p < 0.05, **p < 0.01, ****p < 0.0001 One-way ANOVA.

To explore further the possibility that innate immune activation to mRNA-LNP immunization is altered in GF compared to SPF mice, we extracted RNA from the spleen tissue of mRNA-LNP immunized GF and SPF mice and assessed the expression of a panel of 785 Host Response genes. Principal Component (PC) analysis and unsupervised clustering illustrated that GF and SPF samples clustered separately, primarily along PC2 that explains approximately 17% of variation in the dataset (**Figure 6A**). Pathways associated with myeloid activation, host defense peptides, lysosomes, and phagocytosis were enriched in mRNA-LNP-immunized SPF mice, while pathways associated with lymphocytes, such as TCR and BCR signaling, were increased in GF mice (**Figure 6B**). These data provide further evidence that the innate myeloid response to mRNA-LNP immunization is reduced in GF compared to SPF mice consistent with the flow cytometry analysis. In order to gain insight into what may be driving these differences in myeloid activation we compared the relative expression of genes associated with different innate immune signaling pathways in SPF and GF mRNA-LNP immunized mice. The top signaling pathway that was relatively enriched in SPF samples in this dataset was type-I interferon signaling, while pathways of immune sensing, such as TLR or NLR signaling, where comparatively enriched in GF settings (**Figure 6C**).

**Figure 6:**
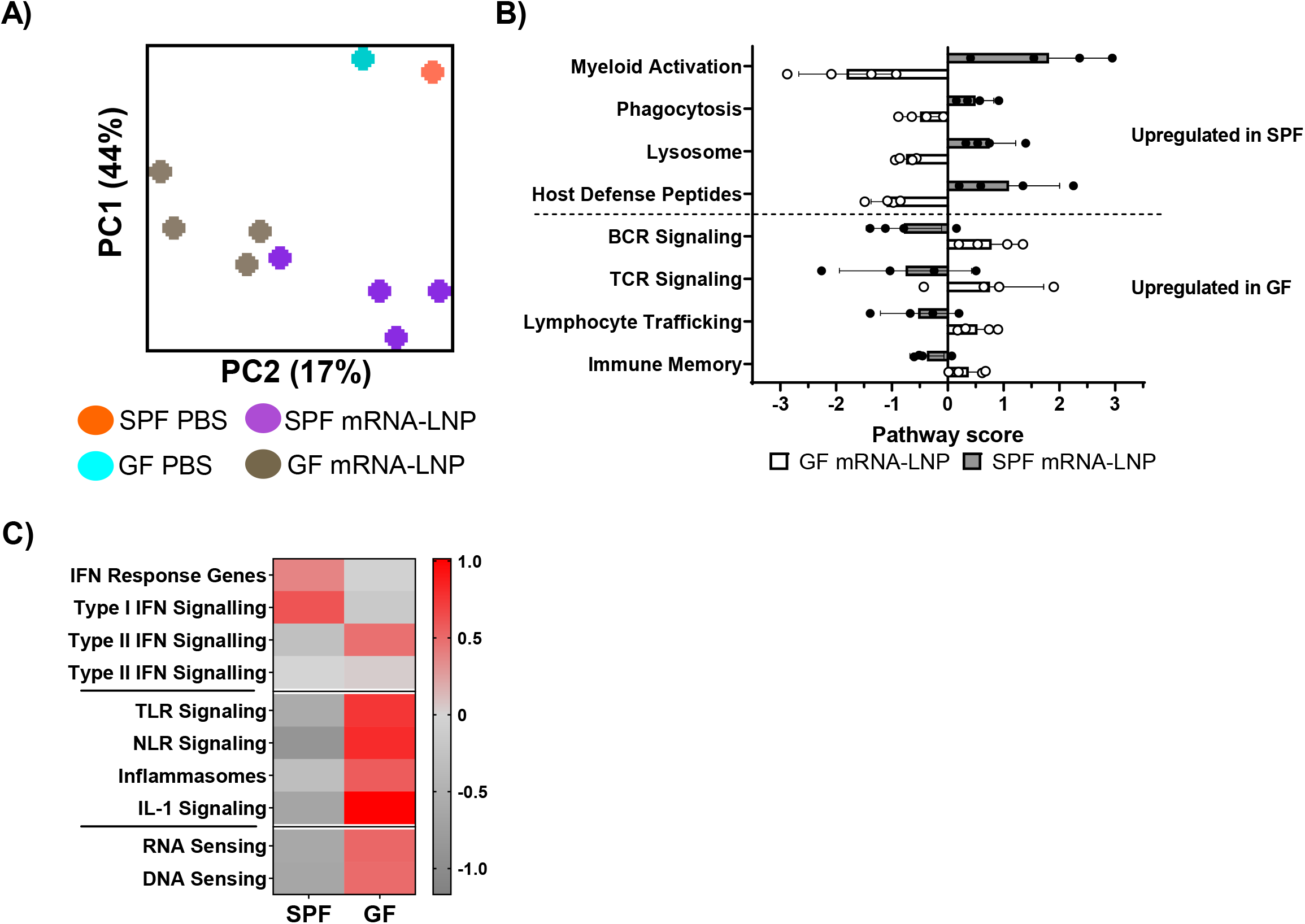
Gene expression profiling of innate immune responses to mRNA-LNP immunization in GF and SPF mice. **(A)** Principle Component (PC) analysis of Nanostring Host Response Gene mRNA Counts from PBS-treated or mRNA-LNP-immunized GF and SPF mice. GF and SPF mRNA-LNP-immunized groups separate along PC2. **(B)** Pathway scores for Myeloid and Lymphocyte pathways in mRNA-LNP-immunized GF and SPF mice. Each data-point represents an individual mouse (n = 4 per group). **(C)** Heatmap representation of innate signaling pathway scores in mRNA-LNP-immunized GF and SPF mice.

## Discussion

We have identified distinct and contrasting ways in which the microbiome impacts DNA and mRNA-LNP vaccine responses. Specifically, the data shown here indicate that the microbiome suppresses humoral and cellular responses to DNA-prime, DNA-Protein-boost vaccination, while the microbiome enhances CD8+ T cell responses to mRNA-LNP immunization. To our knowledge these are the first data demonstrating regulation of these novel vaccine platforms by the microbiome and have implications for our understanding of how differences in vaccine response may be associated with microbiome heterogeneity. Future work will necessarily focus on understanding the full mechanistic basis underlying these different effects.

Our data indicate that the IFN-I response to mRNA-LNP immunization is reduced in GF compared to SPF mice, which is consistent with studies showing that IFN-I at homeostasis and following microbial challenge is microbiome-dependent.^48–54^ The mRNA molecule in this vaccine is nucleoside-modified and stringently purified to eliminate intrinsic immune-stimulatory activity,^55–57^ and therefore we assume the majority of this IFN-I response is being driven by the LNP component. IFN-I is an important regulator of both DNA and mRNA vaccine response, where it’s pleiotropic nature can result in opposing effects on immunogenicity.^58–63^ Prominent effects of IFN-I include enhancing T cell responses through co-stimulation, and recent data have indicated that myeloid activation and CD8+ T cell responses to a nucleoside-modified mRNA-LNP vaccine similar to the one used here are IFN-I dependent.^64^ Consistent with this, GF mice also displayed reduced CD8+ T cell responses and myeloid activation to mRNA-LNP vaccination, and therefore it is likely that an absence of microbiome-dependent IFN-I responses is at least part of the mechanism underlying this effect.

In opposition to such immune-stimulatory IFN-I effects, IFN-I induced by DNA or unmodified mRNA vaccines can also suppress immunogen expression, which inhibits their immunogenicity.^57,60,63,65,66^ We hypothesize that microbiome-primed IFN-I may be suppressing DNA transgene expression in our model providing a potential mechanism for the reduced DNA-prime DNA-protein-boost vaccine responses observed in SPF compared to GF mice. Given that nucleoside-modified mRNA-LNP vaccines are notably less sensitive to IFN-I-mediated suppression,^55–57^ such an outcome would also explain the contrasting contributions of the microbiome to DNA-prime DNA-protein boost and mRNA-LNP vaccine regimens. The low immunogenicity of DNA-HIV-PT123 in mice when delivered s.c. by needle-syringe has prevented us directly testing this hypothesis in our current system. Further work will employ alternative DNA vaccine constructs and delivery methods to alleviate this limitation facilitating further assessment of the sensitivity of DNA vaccines to microbiome regulation and to interrogate underlying IFN-dependent or -independent mechanisms. RNA vaccines that require cytosolic trafficking or amplification or which activate similar IFN-I responses to DNA vaccines may also be sensitive to IFN-I mediated suppression and therefore it will be informative to extend these studies to include unmodified and self-amplifying mRNA vaccine platforms.^55–57,65–68^

Further work is required to understand the innate immune responses elicited by DNA or mRNA-LNP immunization and detail the precise mechanisms that elicit humoral and cellular immunity. Our data report dramatic monocyte activation occurring within 24 hours of mRNA-LNP immunization, as well as accumulation of Ly6C+ and CD64+ DC/macrophage populations that are likely majority monocyte-derived. This is consistent with recent data reporting myeloid activation within 24 hours of mRNA-LNP vaccination in mouse draining lymph nodes and human peripheral blood.^64,69^ The data shown here indicate that this mRNA-LNP-induced myeloid cell activation is regulated by the microbiome warranting further investigation and validation in independent cohorts. This is consistent with profound regulation of mononuclear phagocyte activation by the microbiome in settings of viral infection, which is at least in part IFN-mediated and epigenetically imprinted.^48,49^ T Follicular Helper cell (TFH), Germinal Center B cell, and CD8+ T cell responses to nucleoside-modified mRNA-LNP immunization are independent of Toll-like receptor (TLR)-2, −3, −4, −5, −7 and the TLR adaptor protein Myeloid differentiation primary response 88 (MyD88). TFH and B cell responses are also independent of RIG-I/MDA-5 adaptor protein Mitochondrial anti-viral signaling (MAVS), but are dependent on interleukin 6.^40^ Meanwhile, CD8+ T cell responses to mRNA-LNP are at least partially MDA-5-dependent.^64^ A myriad of innate immune pathways are capable of sensing microbial ligands, exogenous nucleic acid and/or LNP components,^70^ including TLRs (e.g. TLR9), and cytosolic sensors, such as cGAS-STING, RIG-I and AIM2, although the degree to which these various receptors are influenced by the microbiome remains to be determined.

In contrast to our work with nucleic acid vaccines, our studies with protein-adjuvant combinations show that the endogenous mouse microbiome is not necessary for eliciting immune responses towards subcutaneously administered OVA vaccines adjuvanted with Alum, Addavax or AS01, or gp120 protein administered with MF59. This is consistent with published data investigating the contribution of the microbiome to systemically administered Alum- or Cholera Toxin-adjuvanted vaccines in broad-spectrum antibiotic-treated mice.^23,29^ Published data have suggested that the microbiome is necessary for immune responses to OVA adjuvanted with Complete Freud’s adjuvant and to other protein-based vaccines delivered mucosally.^26,29^ Therefore, collectively this work emphasizes that the contribution of the microbiome to protein-based vaccine responses is likely dependent on the nature of the adjuvant as well as the route of delivery. Recent work has also highlighted the microbiome as a potential source of antigen that cross-reacts with pathogen-, tumor- or self-derived antigen.^71–81^ The presence of cross-reactive antigens has been hypothesized to divert HIV vaccine responses away from those that are protective.^80,81^ Future work would be advantaged by extending beyond traditional model antigens, such as OVA, to consider microbiome-vaccine antigen cross-reactivity

As is the case for the majority of microbiome studies in mice, our work is limited to the SPF microbiome of the C57Bl/6 mouse lines sourced as detailed in the materials and methods. By definition SPF microbiomes are constrained to a limited diversity, maintained under careful environmental control, and are exclusive to potentially pathogenic agents. Thus, our work does not exclude vaccine modulatory contributions of microorganisms that are not represented in the mouse colony studied here, but that may be present in the microbiome of other mouse lines or in the human population. Recent data has emphasized the differential responses of environmentally-exposed “Dirty” mice compared to SPF mice following live-attenuated or killed split influenza vaccination.^82^ We completed exploratory vaccine experiments colonizing adult SPF or GF mice with select immunostimulatory species or with WILD-R mouse gastrointestinal material, which increased the diversity of the microbiome community, but was insufficient to modulate the protein-adjuvant vaccine responses tested. We observed that certain taxa of interest, such as WILD-R-derived Proteobacteria, failed to colonize GF mice using the approach described. Future work seeking to further explore the presence of vaccine modulatory microorganisms in the microbiome using gnotobiotic models should extend beyond the C57Bl6/J sources investigated here, and consider approaches that mitigate the limitations of GF colonization (e.g. by colonizing earlier in life).

In conclusion, there is an urgent need to understand the determinants of nucleic acid vaccine immunogenicity and implement this understanding to maximize the potential for clinical success. The work reported here emphasizes the importance of considering the microbiome as a determinant of nucleic acid vaccine immunogenicity. We hypothesize several potential points whereby nucleic acid vaccines may be regulated by the microbiome, including at the level of transgene expression, innate immune response to adjuvant components, and lymphocyte activation/polarization. This establishes a foundation for investigation of microbiome features that may be associated with nucleic acid vaccine response and of candidate mechanisms that may be employed to predict or enhance protective immunity.

## Acknowledgements

We are grateful to Drs. Song Ding and Giuseppe Pantaleo (Eurovacc) for provision of the DNA-HIV-PT123 vaccine, Dr. Alex Maue (Taconic) for provision of WILD-R gastrointestinal contents, and Dr Clarisse Lorin (GSK) for provision of AS01. We are also grateful to the University of Washington Gnotobiotic Animal Core, Fred Hutchinson Cancer Center (FHCC) Comparative Medicine, the HVTN flow cytometry core, and FHCC genomics shared resource for experiment support. This work was supported by the National Institute of Allergy and Infectious Diseases (NIAID; 1R01AI127100-01).

## Author contributions

J.G.K and A.M.F.J conceptualized the study., A.M.F.J., K.H., J.G.K., D.W., M.G.A designed experiments, A.M.F.J., K.H., P.V., N.P., M.G.A. performed experiments and analyzed data. K.M.B., A.F.G, S.M., assisted with experiment design and analyzed data, P.J.C.L. and Y.K.T. provided LNP material. A.M.F.J., and J.G.K., wrote the manuscript.

## STAR Methods

### Mice

SPF C57B6/J mice were obtained from Jackson Labs (Room RB17 or RB05) and housed at Fred Hutchinson Cancer Research Center (FHCRC). GF C57Bl6/J mice were bred in-house at the University of Washington (UW) Gnotobiotic Animal Core (GNAC) and housed in hermetically sealed Tecniplast (West Chester, PA) cages as previously described for the duration of experiments.^83^ Stool was collected from GF mice longitudinally and at the experimental end points and sterility confirmed by Gram staining and sporadic aerobic and anaerobic culture on tryptic soy agar with 5% sheep’s blood and modified yeast casitone fatty acid agar (YCFA; DSMZ 1611). Germ-free breeding colonies are further screened quarterly for presence of bacterial contaminants by amplification of the 16S rRNA gene via PCR. All materials and equipment used with GF animals, such as restrainers for immunization, was sterilized by autoclaving or submersion in 1:3:1 Clidox disinfectant prior to use. Male and female mice were aged 6-16 weeks at the start of immunization regimens. Experimental groups were age-matched and distributed throughout cages during vaccination experiments to reduce confounding influence of ‘cage effects’ due to differences in mouse behavior or SPF microbiome composition between cages. SPF and GF mouse work was approved by the FHCRC and UW Institutional Animal Care and Use Committees, respectively, and all mice were euthanized following AVMA guidelines for CO_2_ overdose.

### Mouse colonization with select taxa or WILD-R gastrointestinal contents

*Akkermansia muciniphila (A. muc, ATC BAA-835), Collinsella aerofaciens (C. aero, ATCC 25986), Fusobacterium varium (F. var, ATCC8501/DSM 19868), Prevotella Copri (P. cop, DSM 18205)* were grown at 37 °C in liquid anaerobic YCFA culture and diluted to OD_600_ 0.5-1. 1 ml of individual cultures were combined, pelleted by centrifugation at 5000 xg, and resuspended in 1ml sterile PBS. 0.5 ml of SFB-containing fecal material (from SFB monocolonized mice) was mixed with the 1-ml bacterial suspension. Each GF mouse received a single 100 μl dose of the pooled consortia via oral gavage. CFUs gavaged per mouse were calculated by serial dilution plating (10^7^-10^8^ for A. muc, C. aero, F. var, and 10^5^-10^6^ for P. cop). Fecal material was collected from colonized mice throughout the experiment, DNA extracted using the QIAamp Fast DNA Stool Mini Kit (Qiagen), and the genome copies of each bacteria quantified by Q-PCR using species specific 16S rDNA primers (see Table 2) relative to standard curves of pure genomic DNA from each strain. Q-PCRs were run using PowerUp SYBR Green Mastermix (Invitrogen) on a Quantstudio 3 thermocycler with the following conditions: 50 °C for 2 minutes, 95 °C for 2 minutes, 35 cycles of 95 °C for 1 second, 60 °C for 30 seconds (adjusted to 55 °C for C. aero), 72 °C for 2 minutes. One week following inoculation mice were immunized with OVA and Addavax as described below. WILD-R and SPF gastrointestinal material from the terminal ileum and cecum was collected anaerobically, weighed, and cryopreserved at a 1:30 dilution (g:ml) in PBS containing 0.1% cysteine and 12%glycerol) as previously described.^84^ GF mice were inoculated with 100 μl of WILD-R or SPF material on three consecutive days. Two weeks following inoculation, blood samples were collected for immune phenotyping using fluorescent-conjugated antibodies (see Table 1) and flow cytometry. Four weeks following inoculation mice were immunized with OVA/Alum as described below. DNA was extracted from residual WILD-R and SPF donor inoculum and from cecal contents of WILD-R-colonized and SPF-C mice using the AllPrep PowerViral DNA/RNA Kit (Qiagen). 16S rDNA amplicon profiling was completed using whoi341 fwd (CCTACGGGNGGCWGCAG) and whoi785 rev (GACTACHVGGGTATCTAATCC) primers with 300bp paired end reads generated on the Illumina MiSeq platform (>20k reads per sample) (www.mrdnalab.com, MR DNA, Shallowater, Tx). OTU tables were generated using the MR DNA ribosomal and functional gene analysis pipeline and alpha and beta diversity metrics calculated and visualized using the tidyMicro pipeline in R.^85^

**Table 1:**
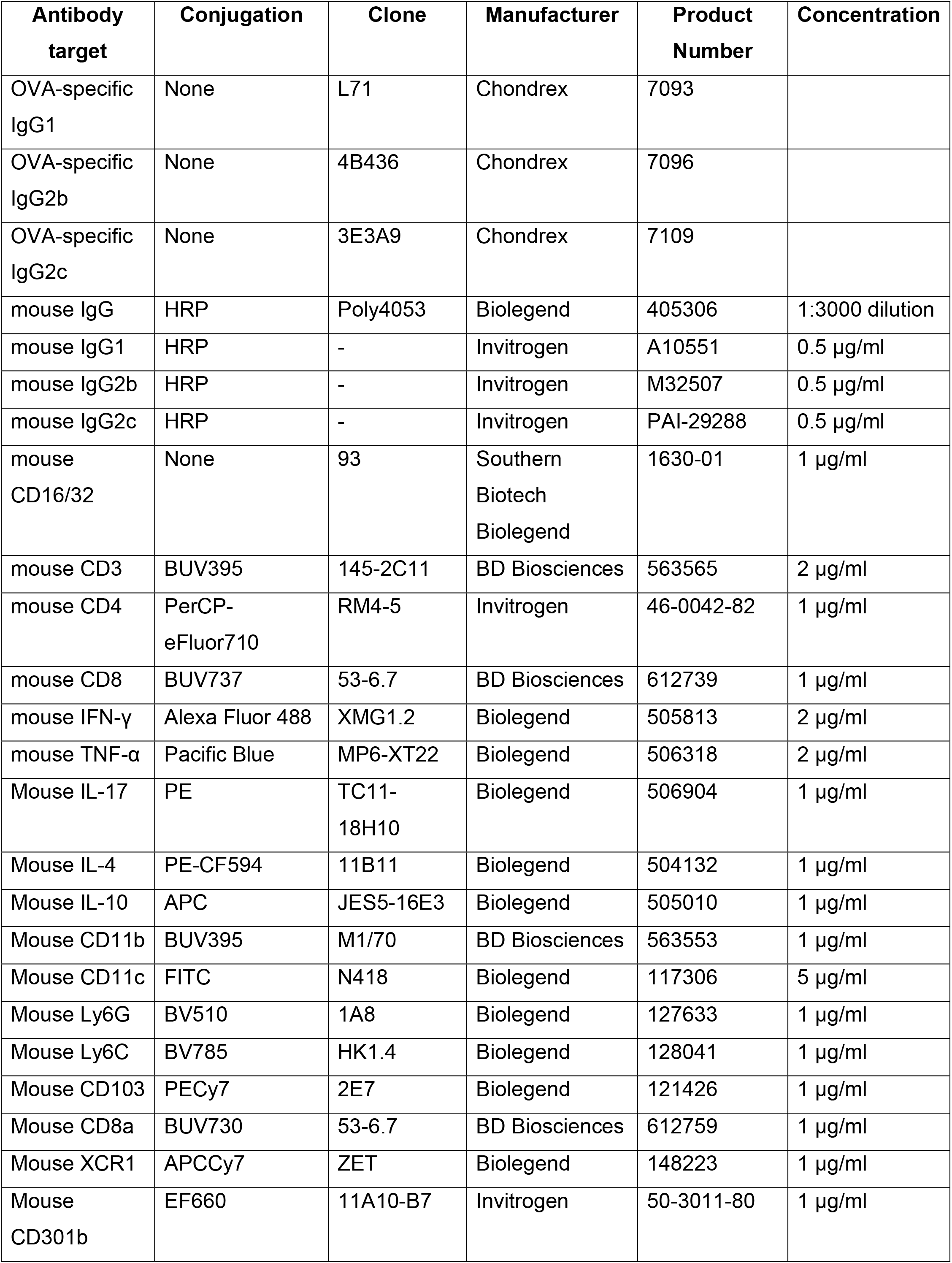
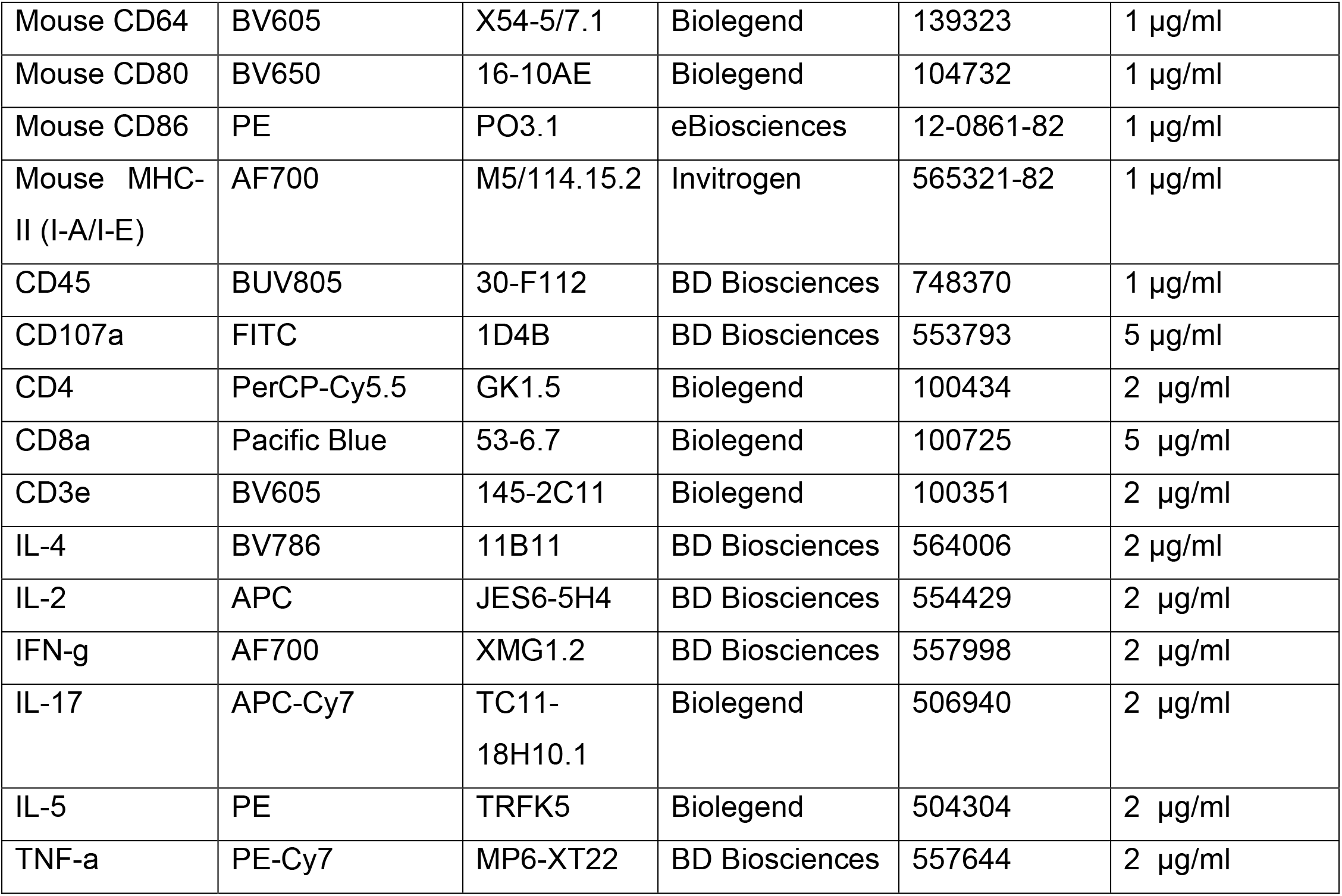
Details of antibody reagents used in these experiments.

**Table 2:**
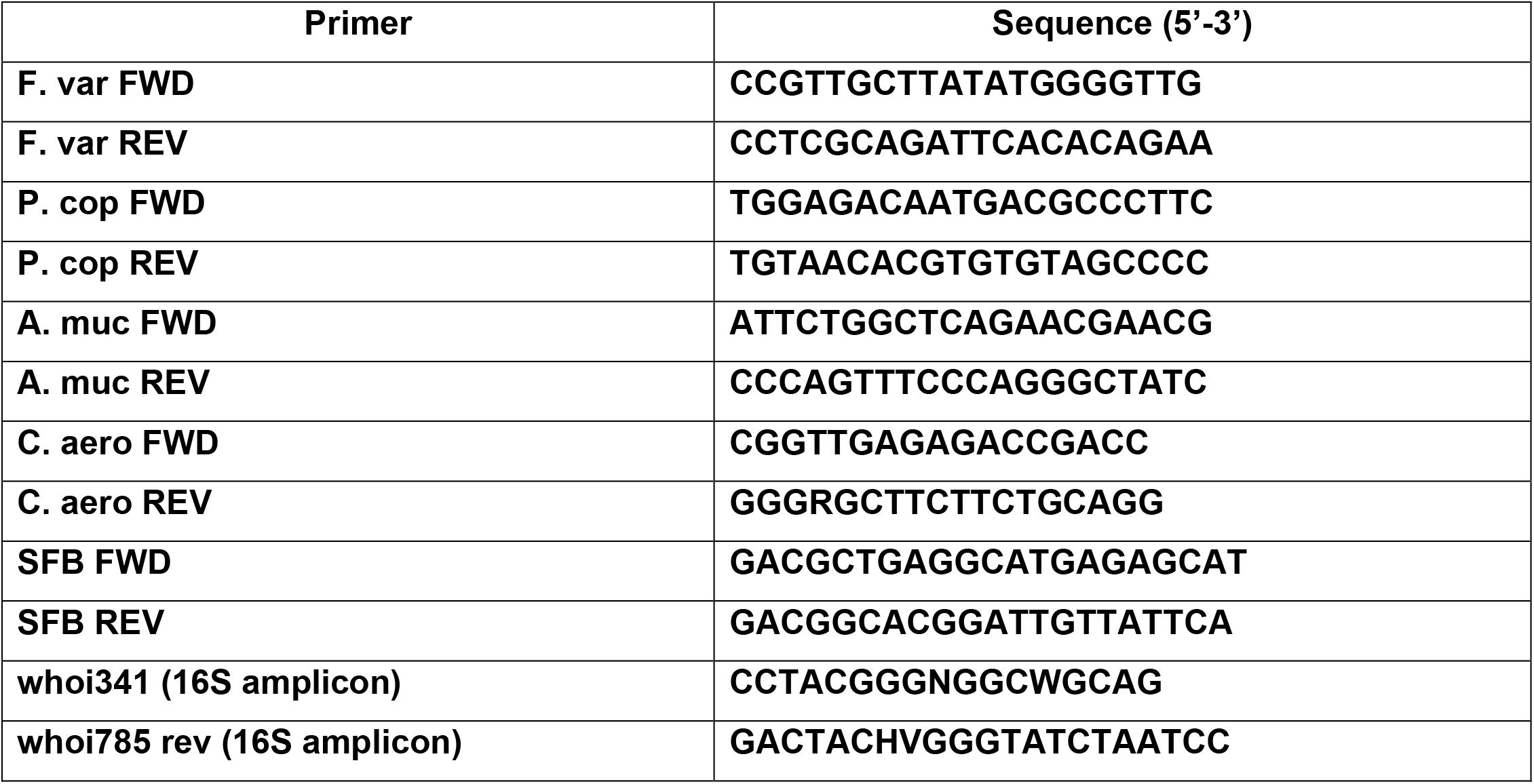
Primer sequences used in these experiments.

### Ovalbumin Protein with Adjuvant Immunization

GF and SPF mice were immunized with a standard prime-boost schema with 10 μg EndoFit Ovalbumin protein (OVA; Invivogen; vac-pova-100) s.c. at the base of the tail (50 μl injection volume) on study day 0 and day 21. OVA was formulated 1:1 with Alhydrogel (Alum, Invivogen; vac-alu-250) or Addavax (Invivogen; vac-1dx-10), or 1:2 with AS01 (kindly provided by GSK), or given unadjuvanted with Ultrapure PBS (VWR; K812-500ML). Negative control mice received ultrapure PBS without OVA by the same route. All vaccine formulations were handled under strict sterile conditions throughout. Mice were euthanized on day 42 and tissue harvested aseptically for subsequent OVA-specific IgG quantification and *ex vivo* OVA re-stimulation assays.

### DNA-HIV-PT123 and gp120 protein immunization

An HIV-1 DNA vaccine, DNA-HIV-PT123, used in HIV Vaccine Trials Network (HVTN) clinical trials HVTN 108 and HVTN111 was provided by Eurovacc for use in these experiments.^44,45^ DNA-HIV-PT123 consists of 3 plasmids expressing (1) HIV-1 Subtype C ZM96 *gag,* (2) HIV-1 Subtype C ZM96 *gp140 ENV,* (3) HIV-1 Subtype C CN54 *pol-nef.* Mice were immunized s.c. at the base of the tail with 100 μg of DNA-HIV-PT123 formulated in ultrapure PBS (VWR; K812-500ML) on study day 0, day 14, and day 35. In select experimental groups, as specified in text and Figures 3 and 4, boost immunizations with 2 μg of bivalent gp120 protein consisting of 1 μg TV.1C gp120 and 1 μg 1086.C gp120 formulated with 1:1 (v:v) MF59 (kindly provided by GSK) or in ultrapure PBS were administered on day 14 and day 35. When delivered on the same day, DNA-HIV-PT123 and bivalent gp120 protein vaccines were prepared separately and delivered contralaterally either side of the tail. Negative control mice received ultrapure PBS by the same route on days 0, 14, and 35. Serum and spleen tissue was collected at day 56 for subsequent gp120- and gag-specific IgG quantification and *ex vivo* gp120 re-stimulation assays.

### mRNA-Spike-LNP Immunization

Codon-optimized coding sequences of Spike receptor binding domain (RBD, amino acids 1-14 fused with amino acids 319-541) of Sars-CoV-2 (Wuhan-Hi-1,GenBank: MN908947.3) were synthesized and cloned into a proprietary mRNA production plasmid. mRNAs were produced to contain 101 nucleotide-long poly(A) tails. m1ψ-5’-triphosphate instead of UTP was used to generate modified nucleoside-containing mRNA. Capping of the *in vitro* transcribed mRNAs was performed co-transcriptionally using the trinucleotide cap1 analog, CleanCap. mRNA was purified by cellulose purification, as described.^86^ All mRNAs were analyzed by agarose gel electrophoresis and were stored frozen at −20 °C. Cellulose-purified m1ψ-uridine containing RNAs were encapsulated in LNPs using a self-assembly process as previously described wherein an ethanolic lipid mixture of ionizable cationic lipid, phosphatidylcholine, cholesterol and polyethylene glycol-lipid was rapidly mixed with an aqueous solution containing mRNA at acidic pH.^87^ The LNP formulation used in this study is proprietary to Acuitas Therapeutics; the proprietary lipid and LNP composition are described in US patent US10,221,127. The RNA-loaded particles were characterized and subsequently stored at −80 °C at an RNA concentration of 1 mg ml^-1^ (in the case of loaded particles) and total lipid concentration of 30 mg ml^-1^. The mean hydrodynamic diameter of mRNA-LNPs was 80 nm with a polydispersity index of 0.02-0.06 and an encapsulation efficiency of 95%. GF and SPF mice were immunized with 1 ug of mRNA-Spike-LNP s.c. at the base-of-the tail in a 100μl volume with ultrapure PBS (VWR; K812-500ML).

### Quantification of Antigen-Specific IgG in Serum

At the experimental endpoints of all experiments, blood was collected by cardiac puncture following euthanasia. Serum was isolated after blood clotted at room temperature for 30 minutes and was centrifuged at 3500 x g for 10 min. All serum was heat inactivated at 56 °C for 30 min with intermittent mixing prior to use in enzyme-linked immunosorbent assays (ELISA) described here.

#### Anti-OVA IgG ELISA

Half area high-binding plates (Corning 3690) were coated by overnight incubation with 200 ng OVA in 0.1 M NaHCO_3_ pH 9.5 at 4 °C. Wells were blocked for 2 hours at 37 °C with 10% nonfat milk (w:v, Research Products International, M17200-500.0) and 0.03% Tween 20 (Fisher Bioreagents, BP337-100) in PBS. Serum was diluted in PBS containing 10% non-fat milk and 0.03% Tween20. OVA-specific mouse IgG1, IgG2b, and IgG2c monoclonal antibodies (Chrondrex) were used to create internal standard curves for quantification of these IgG subclass titers (see Supplemental Table 1 for antibody details). 50μl of diluted serum samples and standard antibodies were added to wells in duplicate and incubated for 1 hour at 37 °C. For detection of anti-mouse IgG1, IgG2b and IgG2c isotypes, HRP-conjugated goat anti-mouse secondary antibodies were added at 0.5 μg/ml in 50 μl PBS containing 10% non-fat milk and 0.03% Tween20 and plates incubated for 1 hour at 37 °C. Plates were developed with 1X Sureblue Reserve TMB (VWR; 95059-290) for 7-15 minutes, depending on isotype, and reactions stopped with 1N H_2_SO_4_ (Fisher Chemicals; SA212-1). Absorbance at 450 nm was read on a SpectraMax i3x plate reader (Molecular Devices) and analyzed using SOFTmax Pro software version 6.5.1. After each incubation step, wells were washed 4 times with 140 μl PBS containing 0.02% Tween20.

#### Anti-gp120 IgG and Anti-Spike IgG ELISA

Half area plates (Corning 3690) were coated overnight at room temperature with 100 ng bivalent gp120 or 100 ng SARS-CoV-2 Spike protein (BEI Resources NR-52397) in 0.1 M NaHCO_3_ pH 9.5. Wells were blocked with 10% nonfat milk (w:v, Research Products International, M17200-500.0) and 0.03% tween 20 (Fisher Bioreagents, BP337-100) in PBS by incubating for 2 hours at 37 °C. Serum was diluted 1:10 in PBS containing 10% nonfat milk and 0.03% Tween20 and loaded onto the plate in duplicate. Subsequently, serum was serially diluted 1:3 across the plate to provide an 11-point dilution curve and plates incubated for 1 hour at 37 °C. Anti-mouse IgG-HRP was added to each well in PBS containing 10% nonfat milk and 0.03 % tween 20 and incubated for 1 hour at 37 °C. Plates were developed and absorbance quantified as described above. Relative anti-gp120 IgG of each sample was calculated as area under the curve (AUC) of a 4-parameter logistic regression of optical density dilution curves.

### *Ex vivo* Splenocyte Antigen Re-stimulation assays

#### OVA and gp120 re-stimulation

Spleens were removed aseptically into sterile cRPMI (RPMI 1640 supplemented with 2 mM L-glutamine (Gibco; 25030081), 10% Fetal Bovine Serum (FBS) (Gibco; 10437028), and 100 U/ml Penicillin-Streptomycin Gibco; 15140122)) and then then disaggregated through a 100 μm mesh filter to generate a single-cell suspension. Cells were washed with 2% FBS in 1x PBS and red blood cells were lysed by incubation in sterile 1x RBC Lysis Buffer (Invitrogen; 501129751) for 3 minutes at room temperature. Cells were washed in 2% FBS in PBS and concentrations of live cells were standardized in cRPMI by manually counting using a haemocytometer. For intracellular cytokine staining (ICS), 1 x 10^6^ splenocytes/well were added to a 96-well round-bottomed plate and incubated in cRPMI alone (unstimulated negative control) or with 10 μg/ml antigen (OVA or bivalent gp120) for 24 hours. After 19 hours, Brefeldin A (5 μg/ml; Biolgend; 420601) and Monensin (2 μM; Biolegend; 420701) were added to block secretion pathways. To quantify secreted cytokines, 5×10^6^ cells/well were added to 12-well plates and incubated with cRPMI alone or 10 μg/ml antigen (OVA or bivalent gp120) in a 2-ml volume for 72 hours. Supernatant was harvested following centrifugation at 400 x g for 5 minutes and stored at −80 °C until analysis. For both ICS and secreted cytokine analyses, positive control wells containing anti-mouse CD3 and CD28 (0.5 μg/ml) were included for each individual mouse. All cell culture incubation was performed at 37 °C with 5% CO_2_.

#### Spike peptide pool re-stimulation

Spleen tissue was collected aseptically, stored in 2-ml complete RPMI and shipped overnight on ice for re-stimulation assays performed in the Weissman lab at University of Pennsylvania. Splenocytes were collected, counted using a ViaCell automated cell counter, and seeded into a 5-mL polypropylene FACS tube at a final concentration of 20,000 cells/μL (100μL/tube). Cells were stimulated with 80μL cRPMI media containing overlapping peptide polls (15-mer, 11 amino acid overlap, 4 amino acid offset, >70% pure as per LC-MS) at a final concentration of 2.5 μl/mL, and 1 μg/mL of anti-CD28/CD49d mixture (co-stimulatory signal) for 6 hours. 1 hour post-stimulation Brefeldin A and Monensin at a final concentration of 5 μg/ml, and 2 μM respectively, was added to block cytokine secretion. All cell culture incubation was performed at 37 °C with 5% CO_2_.

### ICS Flow Cytometry

Following incubation with or without antigen, cells were washed with PBS containing 2% FBS and incubated with Fc Block (anti-mouse CD16/32) and live-dead aqua (1:500 dilution, Invitrogen, L34966) in PBS for 20 minutes on ice. Cells were washed in PBS and then incubated in a surface antibody panel (CD3, CD4, and CD8; see supplemental Table 1) in PBS containing 2% FBS for 30 minutes on ice. Cells were washed with PBS containing 2% FBS, fixed and permeabilized, and stained with an intracellular cytokine panel (IL-4, IL-10, IL-17, IFN-g, and TNF-a) using a Cytofix/Cytoperm ICS kit (BD Biosciences; 554714) according to manufacturer’s instructions. Flow cytometry data for OVA- or gp120-stimulated ICS were acquired on an LSRFortessa X-50 flow cytometer (BD Biosciences) and analyzed on FlowJo version 10.7.1 (FlowJo LLC). Flow cytometry data for Spike-stimulated ICS were acquired on a BD LSR II equipped with 4 laser lines and 18 PMTs. Gates for proportions of cytokine positive cells were set relative to fluorescence-minus-one (FMO) controls containing all antibodies except against the cytokine of interest.

### Quantification of cytokine in antigen-stimulated cell culture supernatant

Concentrations of IL-4, IL-10, and IFN-g were measured in splenocyte culture media supernatant using a Th1/Th2 Mouse Uncoated ELISA Kit (Invitrogen; 88-711-44) following manufacture protocols. Unstimulated and anti-CD3/CD28 samples were included as negative and positive controls, respectively.

### Innate immune response profiling by flow cytometry following mRNA-LNP immunization

24 hours after mRNA-Spike-LNP or PBS s.c. base-of-the-tail administration, spleen tissue was collected aseptically. A small portion of spleen (3 – 5 mm in length) was preserved in RNAlater (Thermofisher) and stored at −80 °C. Remaining spleen tissue was fragmented with a sterile razor blade, placed in 4.5 ml RPMI 1640, 10% FCS, 7.5 mM HEPES with 0.75 mg/ml collagenase II (Sigma-Aldrich) and incubated for 35 minutes at 37 °C with 200 rpm agitation. Collagenase activity was quenched with 200 μl 0.5 M EDTA and tissue disaggregated through a 70 μm cell strainer. Cells were washed with 2% FBS in 1x PBS and red blood cells were lysed by incubation in sterile 1x RBC Lysis Buffer (Invitrogen; 501129751) for 3 minutes at room temperature. Cells were washed in 2% FBS in PBS and concentrations of live cells were standardized by manually counting using a hemocytometer. 2 x 10^6^ cells were incubated with Fc Block (anti-mouse CD16/32) and live-dead Blue (1:500 dilution, Invitrogen, L23105) in PBS for 20 minutes on ice. Cells were washed in PBS and then incubated in a surface antibody panel (supplemental Table 1) in PBS containing 2% FBS for 30 minutes on ice. Cells were fixed (BD Biosciences; 554714) and stored in PBS in the dark at 4 °C overnight prior to data acquisition on BD LSRFortessa X-50 flow cytometer (BD Biosciences) and analysis on FlowJo version 10.7.1 (FlowJo LLC). RNA was extracted from preserved spleen tissue using RNeasy Mini Plus kit (Qiagen), integrity confirmed using Agilent 4200 Tape Station (Agilent) and mRNA transcripts counted in 100ng total RNA using Nanostring nCounter hybridization and quantification protocols. Count data was analyzed using nSolver 4.0 software (Nanostring) with the advanced analysis package installed for pathway scoring.

### Statistics

Prism version 9.0.0 (GraphPad Software, SanDiego, CA) was used to complete statistical analyses. Statistics were calculated for ELISA and ICS results using tests specified in figure legends and were considered statistically significant if *p* ≤ 0.05.

## Supplementary Figure Legends

**Supplementary figure 1:**
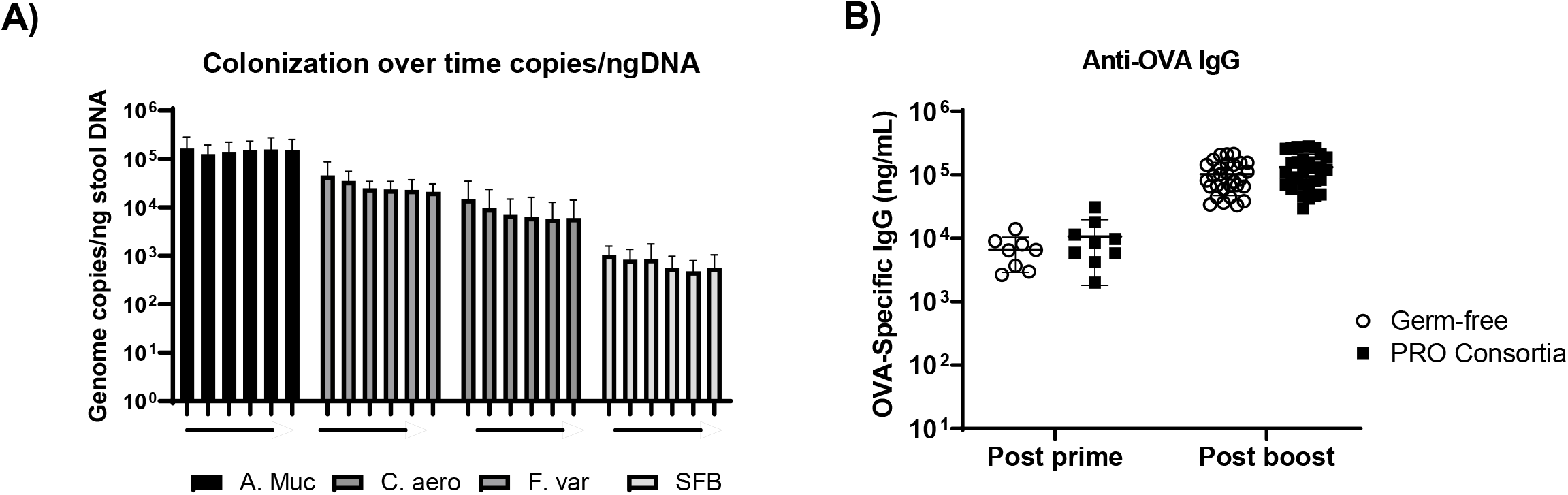
Humoral responses to protein-adjuvant vaccines in selectively colonized mice. **(A)** Genome copies of Akkermansia Mucinophilia (A. Muc), Colinsella Aerofaciens (C. aero), Fusbacterium Varium (F. var), Segmented Filamentous Bacteria (SFB). **(B)** Serum anti-OVA IgG titers of mice colonized with A.Muc, C. aero, F.var, SFB (PRO Consortia) or remaining GF, two weeks post-prime or two weeks post-boost.

**Supplementary figure 2:**
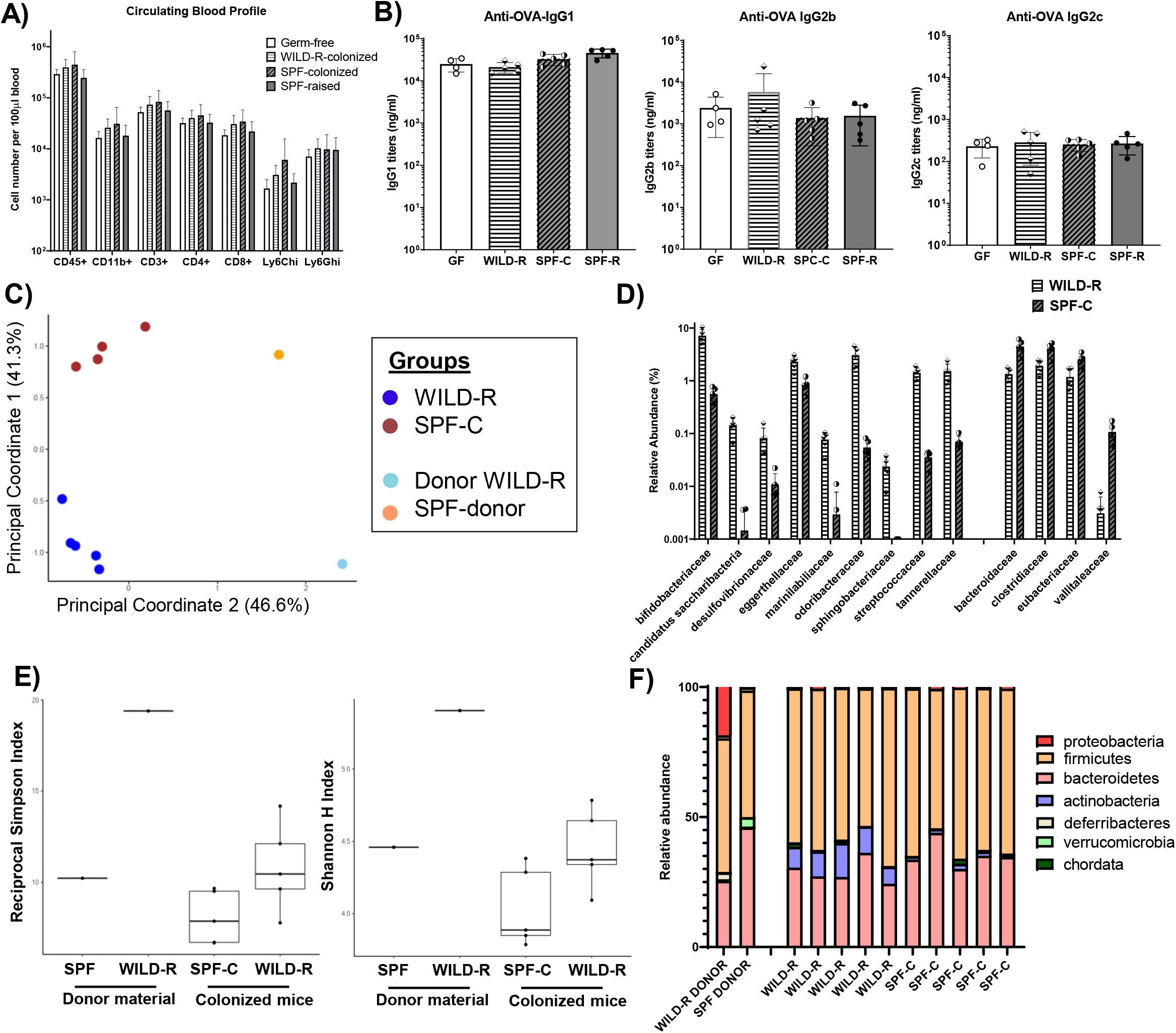
Humoral responses to ovalbumin-adjuvant vaccination in WILD-R or SPF-Colonized mice. **(A)** Numbers of difference cell populations in blood collected two weeks following colonization with WILD-R (n = 5) or SPF (SPF-C; n = 5) material, or of Germ-free (n = 4) and SPF-raised (SPF-R; n = 5) controls quantified by flow cytometry. **(B)** serum anti-OVA IgG1 IgG2b, IgG2c titers two weeks post-boost with OVA/Alum in GF, WILD-R, SPF-C and SPF-R mice. **(C)** Speices-level Principal Coordinate analysis based on Bray-Curtis beta diversity of SPF-C mice (Burgundy dots), WILD-R mice (dark blue dots), or Donor SPF (orange dot) or Donor WILD-R (light blue dot) material. **(D)** Relative abundance of bacterial families significantly different in WILD-R and SPF-C mice based on Multiple T-tests with 1% Benjamini, Krieger, Yekutieli False Discovery Rate correction). Each data-point represents an individual mouse with bars representing mean +/-SD. **(E)** Reciprocal Simpson and Shannon Indices for SPF-C, WILD-R-colonized mice and SPF and WILD-R donor material. Each data-point in A-E represents an individual mouse with bars representing mean +/- SD **(F)** Relative abundance of bacterial Phyla in SPF or WILD-R donor material and in individual WILD-R-colonized (WILD-R C) or SPF-C mice. Each bar represents a single mouse.

**Supplementary figure 3:**
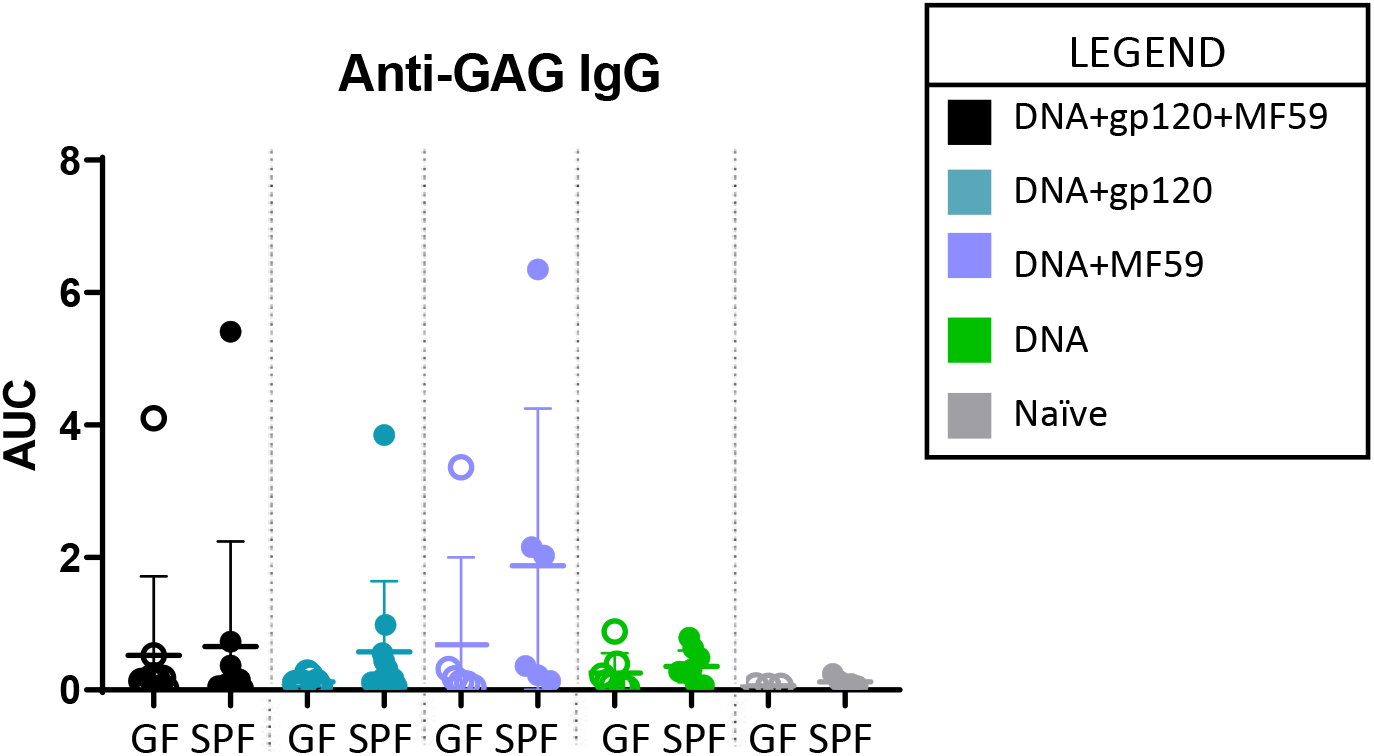
Serum anti-gag IgG in DNA-HIV-PT123 immunized mice. Serum anti-gag IgG titers in GF and SPF mice immunized with DNA-HIV-PT123 (DNA), gp120 and/or MF59 in different combination regimens (as indicated in legend) or unimmunized (naïve) represented as Area Under the Curve (AUC) of an 11-point serial serum dilution.

**Supplementary figure 4:**
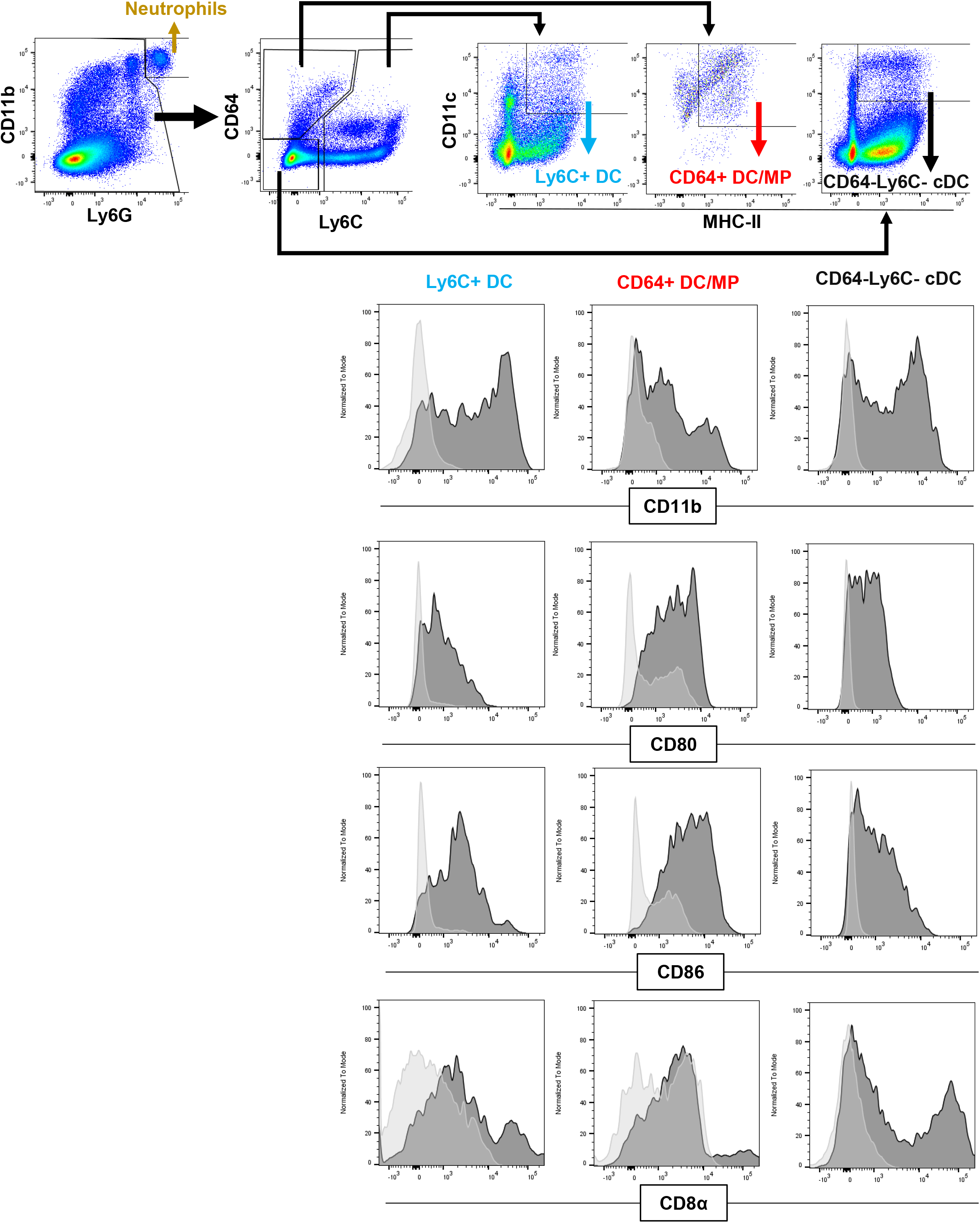
Expression of CD11b, CD80, CD86 and CD8α on Ly6C+ DC, CD64+ DC/MP or CD64-Ly6C-cDC. Representative histograms showing expression of indicated surface protein on splenic Ly6C+ DC, CD64+ DC/MP or CD64-Ly6C-cDC from an unimmunized SPF mouse (dark grey) relative to fluorescent-minus-one (FMO) controls (light grey). Data is representative of n = 4 SPF mice.

**Supplementary Figure 5:**
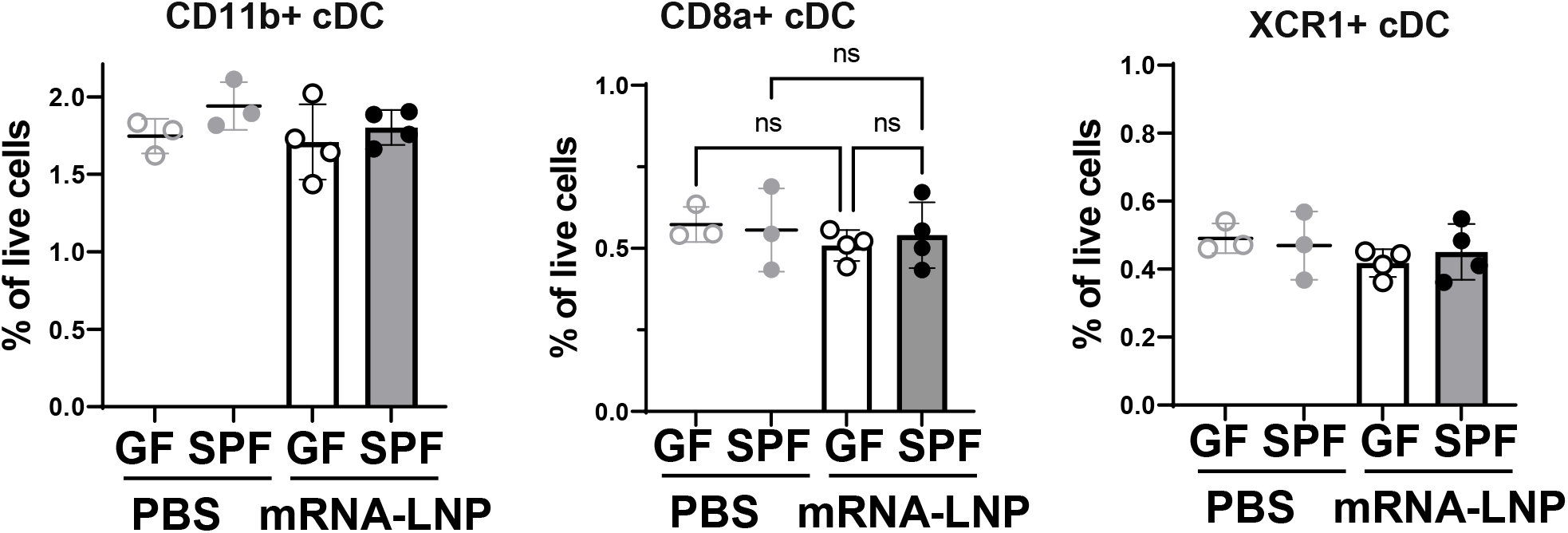
Proportions of CD11b+, CD8a+ and XCR1+ DCs in spleen of GF and SPF mice treated with either PBS or mRNA-Spike-LNP (mRNA-LNP) 24hours post-injection.

